# Does the sequence of a disordered protein encode small molecule binding paths?

**DOI:** 10.64898/2026.05.20.726646

**Authors:** Adelie Louet, Gerhard Hummer, Michele Vendruscolo

## Abstract

Ligand binding to intrinsically disordered proteins resists description in terms of conventional binding pockets, yet it can be analysed as a dynamic process in which ligands move across transient surface interaction sites. Here we characterise a pathway-based representation in which ligand binding is described as a sequence of transitions between residue-defined microstates, enabling ligand-specific effects to be distinguished from intrinsic properties of the peptide conformational ensemble. Using all-atom molecular dynamics simulations of Aβ42 and the C-terminal region of α-synuclein in complex with chemically diverse small molecules, we construct transition matrices that encode ligand movement across the peptide surface and use Markov state models to identify dominant binding pathways and relative binding propensities. Pairwise enrichment-factor and AUC analyses reveal strong conservation of the highest-ranked pathways across chemically diverse ligands, with enrichment factors of 15-45 for the top-ranked states and AUC values typically ≥0.75, well above random expectation. These dominant pathways are also preserved across changes in pH and temperature, whereas a urea control, included as a non-specific binder, shows reduced enrichment, indicating that ligands primarily modulate pathway weights rather than define the underlying network topology. Ensemble docking across chemically diverse libraries further supports the presence of recurrent ligand-accessible hotspots within the peptide conformational ensemble. Building on this framework, we apply a prospective screening pipeline to Aβ42, combining MSM-derived hotspots with sequence-based Ligand-Transformer scoring and Gnina docking across 1.66 million compounds, to nominate 19 candidates for prospective experimental evaluation. Together, these results indicate that disordered protein sequences give rise to conformational ensembles that exhibit characteristic binding pathways for small molecules.

## Introduction

Disordered proteins account for approximately one third of the human proteome and play central roles in signalling, transcriptional regulation, molecular recognition, and the formation of biomolecular condensates^1–4^. Their dysregulation is implicated in a broad spectrum of human diseases, including neurodegenerative disorders associated with the aggregation of Aβ, tau, α-synuclein, and TDP-43^5–8^. Yet disordered proteins remain largely outside the reach of conventional structure-based drug discovery and are still often described as undruggable^5–8^. The difficulty is both conceptual and methodological, since the central organising principle of small-molecule design, the binding pocket, does not straightforwardly apply to proteins that lack persistent three-dimensional structure.

A growing body of experimental and computational evidence shows that small molecules can nevertheless bind disordered proteins^9–16^. Insight into a possible molecular mechanism of binding comes from previous molecular dynamics (MD) studies of folded protein–ligand systems, in which ligand recognition has been reconstructed as a sequence of metastable intermediates and surface-diffusion events rather than direct insertion into a binding pocket^17,18^. An analogous picture has emerged for disordered proteins, where NMR spectroscopy, single-molecule fluorescence microscopy, and long-timescale MD studies indicate that ligands engage disordered proteins through a series of transient contacts rather than through occupancy of a single pose. Fasudil binds the C-terminal region of α-synuclein by shuttling among charge-charge and π-stacking interactions without locking into a fixed geometry^19^, 5-fluoroindole binds the disordered domain of NS5A with sub-mM affinity while retaining picosecond rotational mobility in the bound state^20^, and 10074-G5 (G5) binds monomeric Aβ42 with low-µM affinity while preserving, and even enhancing, conformational entropy^21,22^. In these systems, the bound state remains disordered, as binding emerges from many weak, transient interactions rather than from complementarity to a defined cavity.

This dynamic mode of recognition has been described as fuzzy binding, dynamic shuttling, or disordered binding. We recently formalised this picture by introducing the concept of a binding path, which is a stochastic trajectory through a network of transient residue clusters contacted by a ligand on the surface of a disordered protein^23^. In this framework, small-molecule binding is represented as a Markov state model (MSM) on the space of residue contacts^23^. Each state corresponds to a small set of residues simultaneously contacting the ligand, and transition probabilities describe how the ligand redistributes its contacts over time^23^. Applied to G5 binding to Aβ42 and fasudil binding to α-synuclein, this approach recovered dissociation constants consistent with experiment and showed that ligand motion is dominated by gliding transitions between overlapping residue clusters, with rare long-range hops^23^.

This representation raises a central question. If binding to a disordered protein is not occupancy of a pocket but motion through a network, what determines the network? One possibility is that binding paths are primarily ligand-defined: different ligands carve different routes through the disordered ensemble, so each ligand-protein pair requires its own structural characterisation. A second possibility is that the network is predominantly protein-defined: the sequence, through the conformational ensemble it populates, presents recurrent surface regions that different ligands sample with different weights. The distinction is important for drug discovery. In the first case, each chemical series must be mapped independently. In the second, the druggable features of a disordered protein could be mapped as intrinsic properties of the protein ensemble and then used to guide ligand discovery.

Here we test these alternatives by constructing binding-path MSMs for two disease-relevant disordered systems: Aβ42 and the C-terminal region of α-synuclein. We analyse a panel of chemically diverse small molecules, a urea control, and three environmental conditions for Aβ42-G5, varying pH and temperature. Across these systems, we compare ranked binding pathways using enrichment-factor and AUC analyses, asking whether the highest-probability residue-contact pathways are preserved when the ligand or environment is changed. We then use ensemble docking across chemically diverse compound libraries as an orthogonal computational test of whether high-probability surface regions recur across ligands or are ligand-specific.

Our results indicate that the dominant binding pathways and surface hotspots of these disordered peptides are largely properties of the peptide conformational ensemble rather than of any single ligand. Chemically diverse ligands preferentially visit similar regions of the sequence, pathway conservation is preserved across changes in pH and temperature, the urea control departs from this pattern, and ensemble docking identifies recurrent ligand-accessible regions. Building on the finding that hotspots are protein-defined, we apply a prospective Aβ42 screening pipeline that combines MSM-derived hotspots, sequence-based Ligand-Transformer scoring^24^, and Gnina docking^25^ to nominate nineteen candidate ligands from a 1.66-million-compound library for experimental evaluation.

Together, these results extend the binding-path framework from a description of how individual ligands engage disordered proteins to a comparative framework for identifying recurrent binding routes within disordered conformational ensembles. In drug discovery, this shifts the relevant design problem from fitting a ligand into a static pocket to matching ligands to dynamic binding paths defined by the disordered protein ensemble, which is in turn encoded in the sequence of protein.

## Results

### A transition-based representation of binding to disordered peptides

To investigate small-molecule binding in disordered protein systems, we analysed two targets and a panel of chemically diverse ligands (**Figure 1**). For Aβ42, three ligands were examined: (i) the previously reported G5^21,22^, and D4 and D8, identified through Ligan-Transformer screening^26^, and (ii) for the C-terminal region of α-synuclein, seven inhibitors were selected from previously reported molecular dynamics trajectories^19^. Aβ42-G5 was additionally simulated under three conditions: pH 5 at 278 K, pH 7 at 278 K, and pH 5 at 298 K, to test the sensitivity of binding behaviour to physiological perturbation.

**Figure 1.**
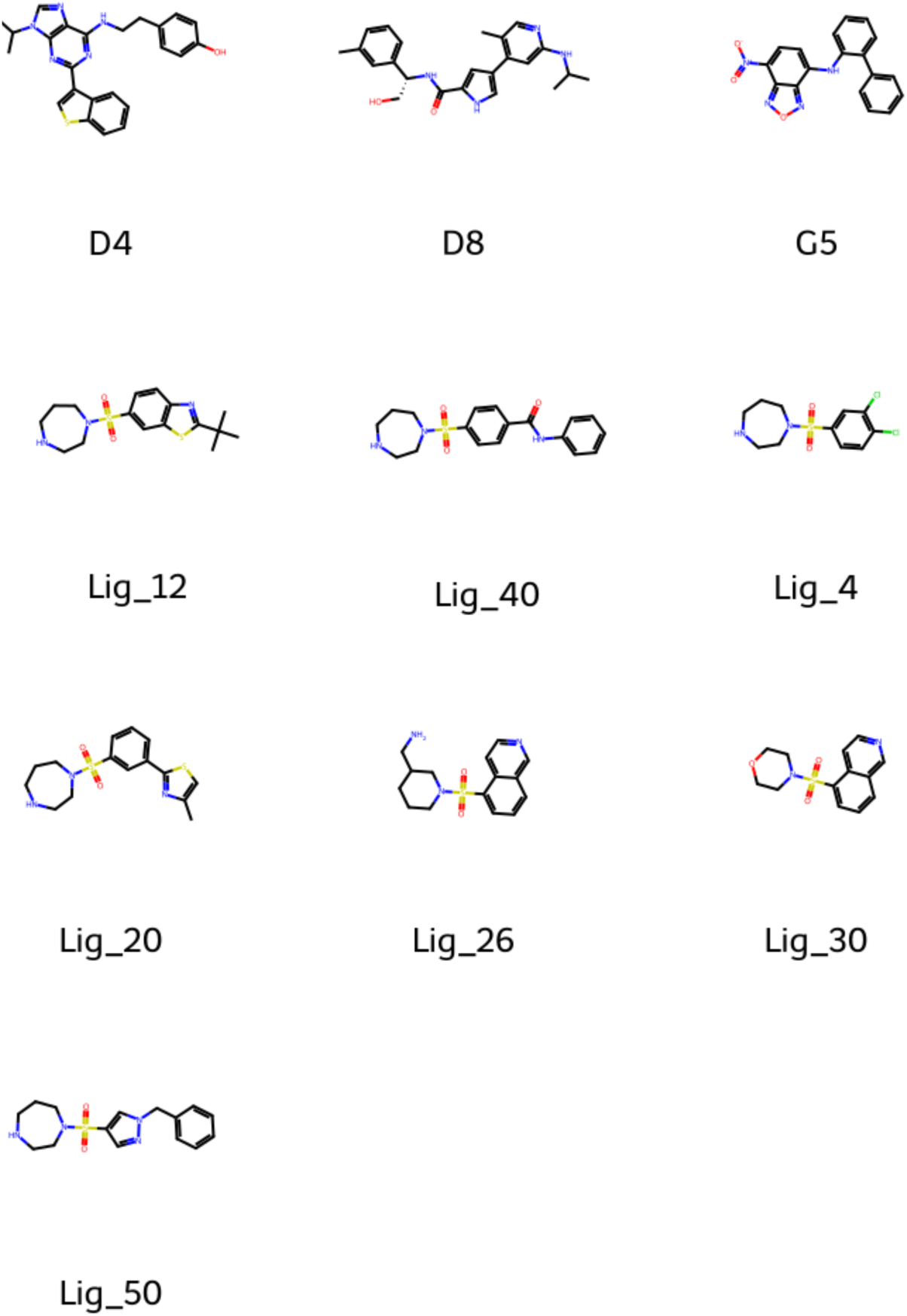
Chemical structures of the small molecules used to characterise binding pathways across three disordered systems. Top row: ligands studied in complex with Aβ42 (D4 and D8), identified through computational screening, and G5 (10074-G5), a previously characterised inhibitor of Aβ42 aggregation. Second row: seven ligands (Lig_4, Lig_12, Lig_20, Lig_26, Lig_30, Lig_40, Lig_50) selected from previously reported molecular dynamics trajectories of the C-terminal region of α-synuclein. The set spans a wide range of molecular sizes, scaffolds, and physicochemical properties, providing the chemical diversity required to test whether dominant binding pathways are ligand-specific or are properties of the peptide conformational ensemble.

A natural starting point for analysing these complexes is the time-averaged residue-ligand contact frequency, which summarises how often each residue interacts with the ligand over the trajectory (**Figure 2a**). Such static profiles are useful for ranking residue involvement, but they average over bound, transiently bound, and unbound configurations and therefore obscure the dynamic character of the interaction. In both systems, ligands were observed to diffuse along the peptide surface rather than occupy a single pose: G5, for example, traverses the surface of Aβ42 forming transient interactions with multiple side chains, consistent with previous reports. Capturing this behaviour requires a representation that tracks how ligand-residue contacts evolve in time, not only which residues are contacted on average.

**Figure 2.**
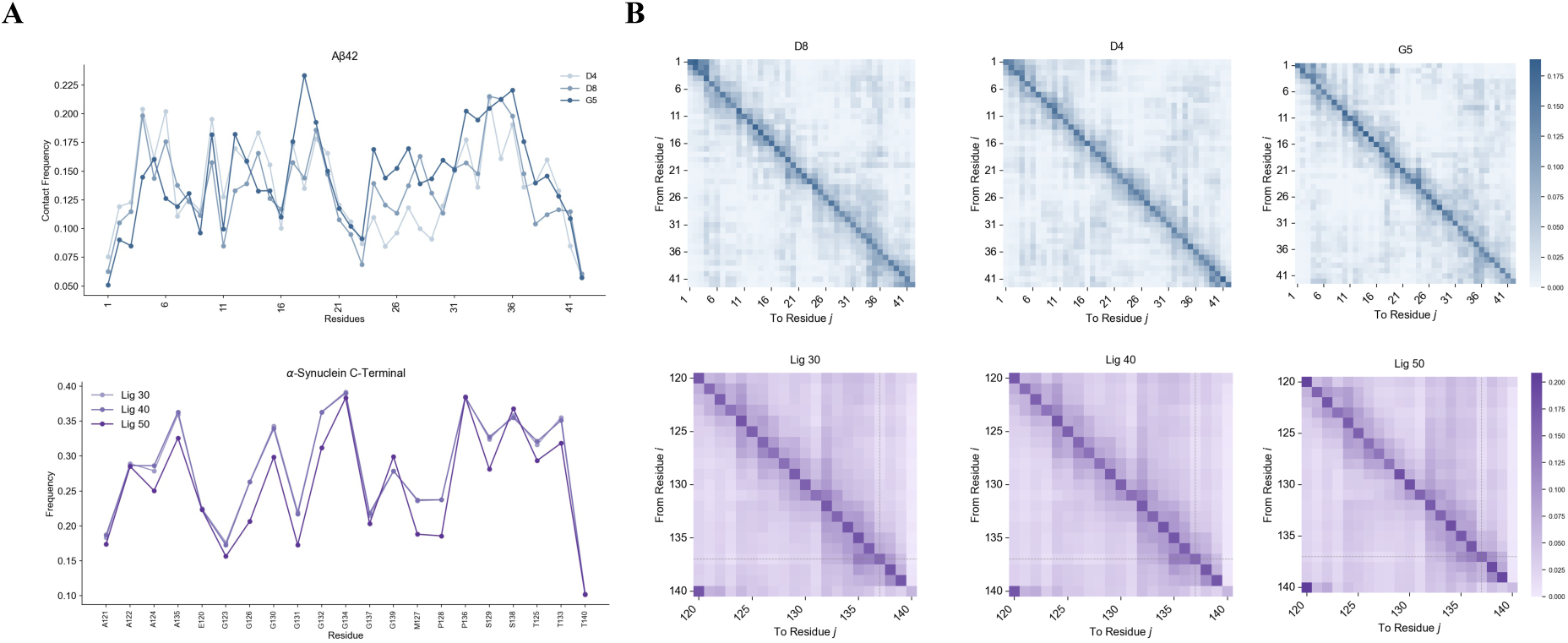
Static and dynamic representations of ligand binding across the two disordered systems. **(a)** Static binding profiles for Aβ42 (top, blue), and the C-terminal region of α-synuclein (bottom, purple), shown as normalised residue–ligand interaction frequencies along the peptide sequence. Each line corresponds to a distinct ligand, with profiles for chemically diverse ligands within each system tracking closely and identifying shared regions of preferential contact. **(b)** Handoff matrices encoding the dynamic redistribution of ligand contacts across the peptide surface. Each matrix element H_ij_ records the number of transitions in which the ligand contacts residue i at frame t and residue j at frame t+1, irrespective of whether the contact with residue i is retained. Matrices are shown for the three Aβ42 ligands (D8, D4, G5; top row), and three representative α-synuclein C-terminal ligands (Lig_30, Lig_40, Lig_50; bottom row). The strong diagonals indicate persistence of contacts across consecutive frames, while off-diagonal features identify the residue pairs between which the ligand most frequently transfers. Together, panels (a) and (b) motivate the transition from static interaction frequencies to a pathway-based description in which ligand binding is represented as motion through a network of residue-defined microstates.

We therefore constructed a time-resolved contact representation in which a transition is recorded whenever residue i contacts the ligand at time t and residue j at time t+1, regardless of whether contact with residue i persists. Summing these events over each trajectory yields a handoff matrix that captures the redistribution of ligand contacts across the peptide surface (**Figure 2b**). The handoff matrices reveal distinct binding hotspots and structured off-diagonal patterns rather than uniform diffusion, indicating that ligand motion is dominated by a relatively small set of preferred transitions.

To formalise this representation, we built Markov state models (MSMs) on the space of residue contacts. Each microstate is defined as a quintuplet of residues simultaneously contacting the ligand, and transitions between microstates encode the redistribution of contacts at each simulation step. The transition matrix obtained from this construction can be visualised as a knowledge graph (**Figure 3a**), in which nodes correspond to residue-contact microstates and edges to inter-state transitions, with node size proportional to degree centrality.

**Figure 3.**
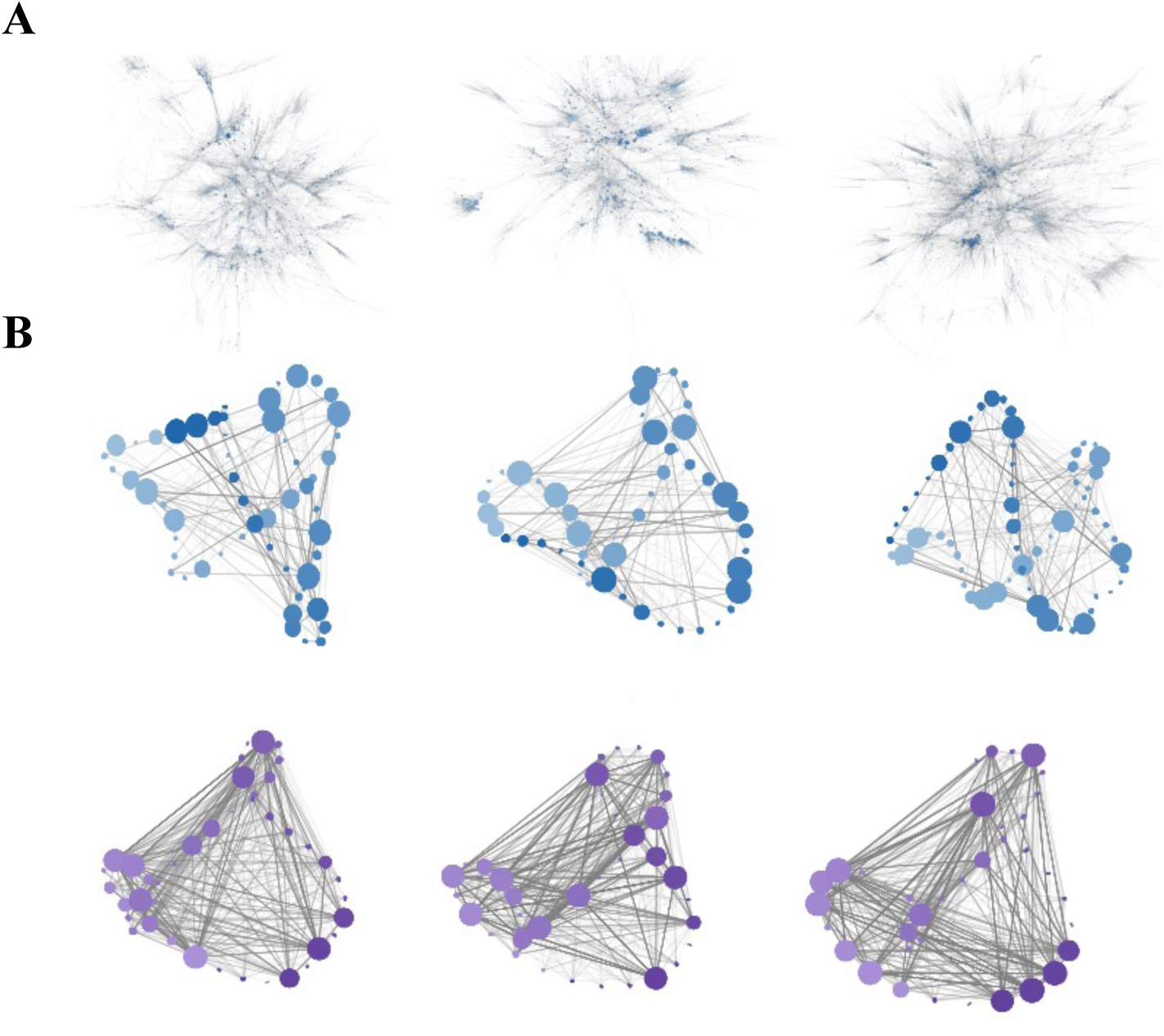
Markov state model representation of binding pathways. **(a)** Microstate networks for Aβ42 (top), and α-synuclein C-terminal (bottom), with each node representing a quintuplet of residues simultaneously contacting the ligand and edges representing observed transitions between microstates. Node size is proportional to degree centrality. The dense, interconnected topology of each network reflects the diffuse binding landscape characteristic of disordered systems. **(b)** Macrostate networks obtained by aggregating microstates within each Louvain community into a single representative node, providing a coarse-grained view of the binding landscape. Node size reflects the summed equilibrium population of the community, and edges represent inter-community transition flux. High-affinity macrostates were identified on the basis of their dissociation constants K_D_, derived from the equilibrium distribution of the MSM.

Binding equilibria were obtained by solving the master equation at steady state, following our previous work. From the equilibrium distribution *P*_eq_, a relative dissociation constant was assigned to each binding state as *K_D,i_* = *p_u_*_,eq_/*p_i_*_,eq_, where *p_u_*_,eq_ is the equilibrium population of the unbound state and *p_i_*_,eq_ the population of state i. To enable comparison across systems with different total bound populations, these relative values were converted into effective equilibrium dissociation constants using the simulation box concentration of the ligand, reported below in μM. Microstates were further clustered into macrostates using network-based aggregation, revealing groups of frequently interconverting states that define dominant binding pathways (**Figure 3b**), and representative bound configurations within each macrostate could be extracted directly from the trajectory (**Figure 3c**).

This framework enabled quantitative ranking of binding affinities across systems. At a 5.5 Å contact cutoff, the effective KD for the α-synuclein C-terminal complex was 8.89 × 10^3^ μM, consistent with the highly transient surface engagement expected for this disordered peptide segment (ligand-bound occupancy ∼45%). Aβ42 ligands gave markedly tighter values, spanning 4.32 × 10^2^ to 1.08 × 10^3^ μM across the G5, D8, and D4 compounds, in line with their more persistent binding (occupancy 71-81%) and with the low-μM affinities previously reported experimentally for G5; the computed values reflect the global KD averaged over all binding configurations rather than a single optimized pose.

### Binding path independence from ligands

To determine whether the binding transition networks identified above are dictated by ligand identity or instead reflect intrinsic properties of the peptide systems, we quantitatively compared ranked transition pathways across all ligand-bound conditions (**Figure 4**). For each system, the residue-contact triplets identified by the MSM were ranked by equilibrium probability, and pairs of ranked lists were compared using enrichment factor and AUC analyses.

**Figure 4.**
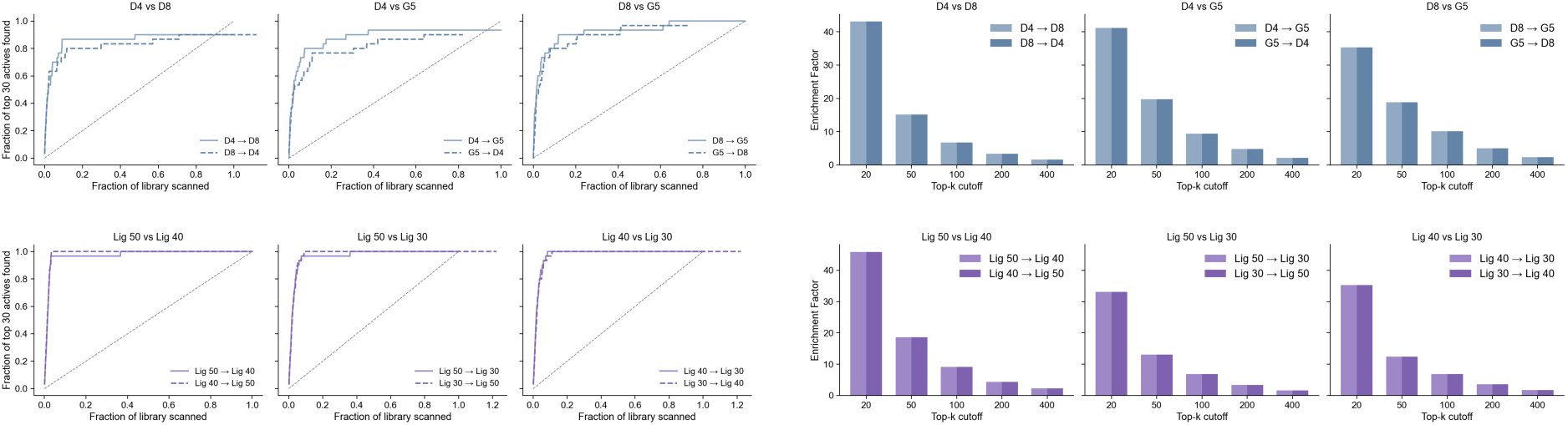
Conservation of dominant binding pathways across chemically diverse ligands. Pairwise cross-ligand comparison of ranked binding pathways for the three Aβ42 ligands (top row, blue), and three representative α-synuclein C-terminal ligands (bottom row, purple). **(a)** Cumulative recovery curves showing the fraction of top 30 reference states (defined as the highest-probability residue-contact triplets in one system) recovered as the ranked list of the comparison system is scanned from highest to lowest probability. The diagonal grey line indicates random expectation. Across all ligand pairs except those involving the urea control, recovery rises rapidly above the random baseline, with the majority of reference states recovered within the first 20-30% of the ranked list (AUC values typically ≥0.75). **(b)** Pairwise enrichment factors for overlap of top-ranked contact triplets at top-k cutoffs of 20, 50, 100, 200, and 400. Enrichment factors of 15-45 are observed at the smallest cutoffs for chemically related ligand pairs, decaying smoothly with increasing k but remaining above unity at large cutoffs. The urea control departs from this pattern, with markedly reduced enrichment relative to the structured ligands, as expected for a non-specific binder. Together, the recovery curves and enrichment factors indicate that the dominant binding pathways are largely ligand-independent: chemically diverse ligands sample the same high-probability residue-contact pathways, with ligand identity modulating pathway weights rather than redrawing the underlying transition network.

Pairwise enrichment factors revealed strong and systematic agreement among the highest-ranked binding states (**Figure 4a**). For the smallest cutoffs (top 20-50 states), enrichment factors were consistently well above random expectation, with EF of 15-45 across pairs of specific binders within each system. Enrichment decreased smoothly with increasing cutoff size but remained above unity even at large cutoffs (top 200-400), indicating that agreement extends well beyond the most probable transitions. This conclusion is reinforced by AUC enrichment curves (**Figure 4b**), in which top-ranked states from one system were treated as reference actives and their recovery was monitored as the ranked list of a second system was scanned from highest to lowest probability. Across all pairwise comparisons of specific binders, AUC values were typically ≥0.75, well above the random baseline of 0.5, and in most cases a large fraction of the dominant pathways was recovered after scanning only 20-30% of the ranked list. Minor directional asymmetries were observed, reflecting differences in the ordering of lower-probability states, but the overall enrichment behaviour was conserved across all pairs of specific binders.

We further tested whether this conservation extends to environmental perturbation by comparing Aβ42-G5 binding pathways across three physiological conditions: pH 5 at 278 K, pH 5 at 298 K, and pH 7 at 278 K (**Figure S1**). Across all three pairwise comparisons, we observed strong enrichment of the highest-ranked states, with EF values comparable to or exceeding those seen between different ligands on the same target. The dominant binding pathways are therefore conserved not only across chemically diverse ligands but also across changes in pH and temperature, indicating that the underlying transition network is a robust property of the peptide conformational ensemble rather than a feature of any particular simulation condition.

Together, these results indicate that dominant binding pathways are largely ligand-independent. Ligands modulate the weights and residence times along an existing transition network without substantially redrawing its topology, and a non-specific binder is distinguishable from specific binders in the same analysis.

### Ligand-independent identification of binding hotspots by ensemble docking

To test whether the binding regions identified by MSM analysis correspond to genuine ligand-accessible surface features, and whether their identification depends on the chemistry of the bound ligand, we applied an ensemble docking strategy across all systems. For each holo trajectory, a random subset of representative frames was selected, and candidate pockets were identified on each frame using Fpocket. Each pocket was then evaluated by docking a chemically diverse subset of the Lipinski-filtered ZINC15 library (constructed as described in Methods) with AutoDock Vina, yielding a distribution of docking scores for every pocket across the ensemble. This procedure decouples pocket identification from any single ligand structure and provides a high-throughput readout of pocket quality.

Across all three systems, docking scores were strongly pocket-dependent rather than uniform (**Figure 5**). For each target, a subset of pockets consistently yielded favourable Vina scores (typically −7 to −8 kcal/mol) across the chemically diverse library, whereas other pockets scored poorly (−3 to −5 kcal/mol) regardless of ligand chemistry. The line-by-line traces in Figure 5, each corresponding to a single docked compound, show that this stratification is largely conserved across compounds within a system, as ligands of different chemotypes agree on which pockets are favourable and which are not. This is the docking-side counterpart of the MSM result, where a small number of recurrent surface regions is preferentially sampled across chemically distinct probes.

**Figure 5.**
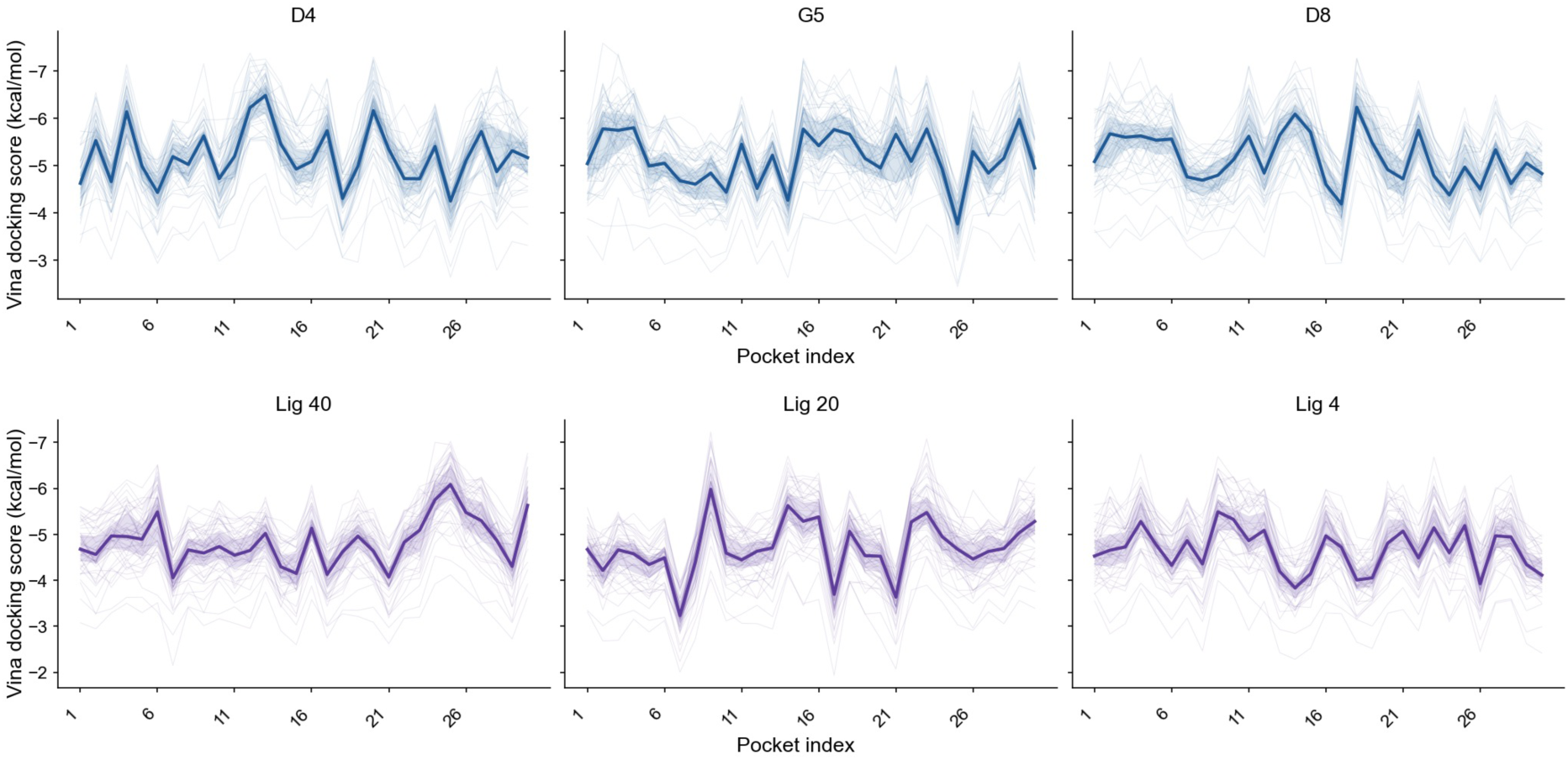
Ligand-pocket docking score landscapes across the two disordered systems. AutoDock Vina docking scores for individual ligands across the top 30 binding pockets identified for each system, with one panel per ligand. Each line corresponds to a distinct docked pose, coloured by ligand identity within a system: Aβ42 ligands D4, G5, and D8 (top row, blue); and α-synuclein C-terminal ligands Lig_40, Lig_20, and Lig_4 (bottom row, purple). Pocket index is shown on the x-axis and Vina docking score (kcal/mol, lower values indicate stronger predicted binding) on the y-axis. Within each panel, distinct pocket-dependent score patterns are observed rather than uniform scores across pockets, indicating structured binding preferences. Across ligands within a system, the high- and low-scoring pockets are largely shared: pockets that dock well for one ligand tend to dock well for chemically distinct ligands, and pockets that dock poorly do so consistently. This pattern indicates that productive binding pockets are intrinsic features of the peptide conformational ensemble and can be identified without reliance on any single ligand chemistry.

To assess whether these recurrent docking sites correspond to the binding regions identified from the dynamics, we compared the MSM-derived hotspots (the highest-affinity macrostates) with the top-scoring docking pockets from the same trajectory ensemble (**Figure S2**). Ensemble docking recovered the dynamics-defined hotspot in 89% of cases, compared with 26% for Fpocket detection alone on the same set of frames. The two analyses, which use entirely different definitions of a binding region (high equilibrium population of a residue-contact state versus favourable docking score against a diverse compound library) therefore converge on the same surface regions.

Taken together, the docking analysis indicates that productive binding regions are intrinsic features of the peptide conformational ensemble rather than properties of any single bound ligand, and that they can be recovered by an entirely structure-based readout that does not use the bound ligand for pocket detection.

### Prospective Application to Aβ42

To test the translational value of the framework, we applied it prospectively to Aβ42 with the aim of identifying small-molecule ligands of greater overall binding affinity than previously reported binders. The pipeline follows directly from the protein-defined picture developed above: if binding hotspots are intrinsic to the conformational ensemble, they can be mapped once and then used to prioritise chemistry, rather than re-derived independently for every chemical series. We therefore implemented two parallel filtering steps, one for hotspot identification and one for ligand prioritisation, before combining them in a hotspot-by-ligand docking screen.

For hotspot identification, we used the Aβ42-G5 MSM to extract the 40 highest-affinity binding microstates, ranked by equilibrium population. The representative frame of each microstate, with G5 removed, was retained as an apo receptor structure, yielding a panel of 40 conformations of the Aβ42 ensemble enriched in surface features identified by the binding-path analysis. For ligand prioritisation, the TargetMol isomeric library (∼2 × 10^6^ compounds) was scored against the Aβ42 amino acid sequence using Ligand-Transformer, a sequence-based affinity predictor that does not require a three-dimensional receptor and is therefore suited to disordered targets. The 20,000 compounds with the highest predicted pK_d_ for the full Aβ42 sequence were retained for structure-based screening.

Each of these 20,000 compounds was then docked independently into every one of the 40 hotspot conformations using Gnina, and ranked by mean CNN affinity score across all pockets. The top 200 compounds (**Figure 6**) showed mean CNN affinity scores in the range 7.0-7.6 (pK_d_ units), with the highest-ranked compound reaching a mean score above 7.6. By comparison, G5 (the highest-affinity Aβ42 binder previously characterised in our group) falls in the low-μM regime experimentally, corresponding to pK_d_ of 6, indicating that the predicted affinities of the top hits exceed those of G5 by approximately one order of magnitude on the CNN scale. The substantial error bars in Figure 6 reflect variability across the 40 hotspots, consistent with the picture that individual ligands engage some hotspots more favourably than others while remaining competitive across the panel.

**Figure 6.**
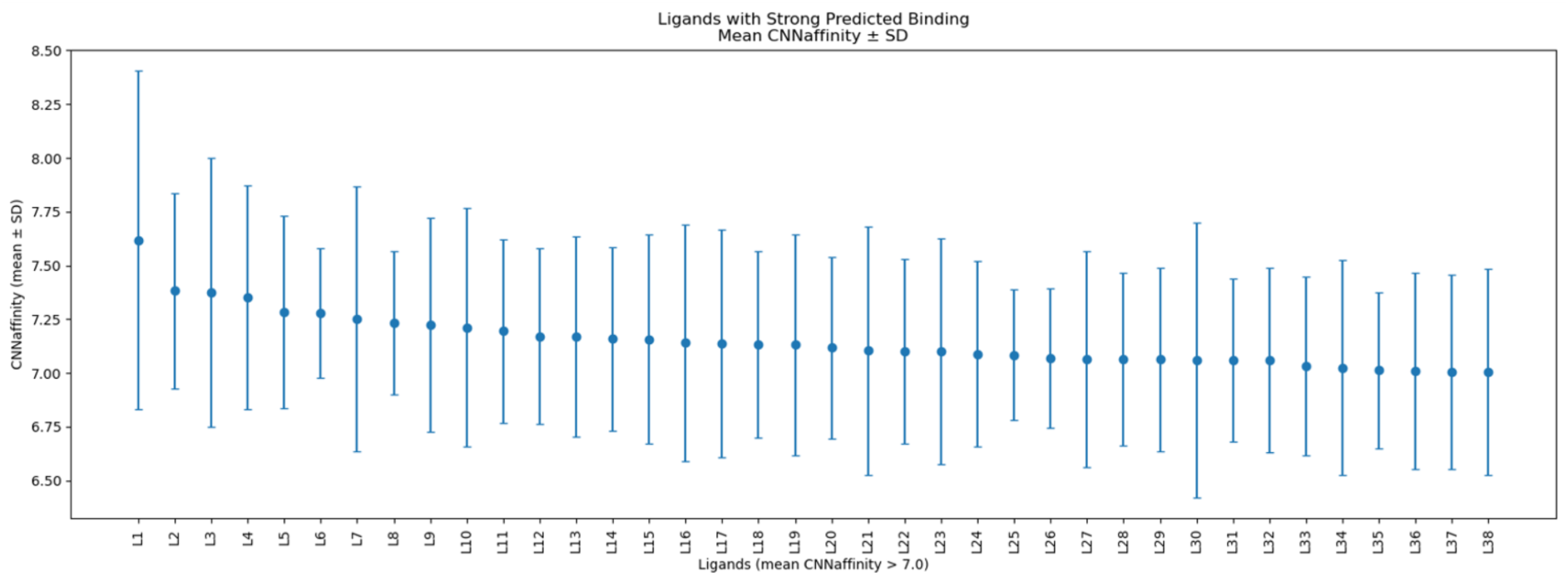
Prospective ranking of candidate Aβ42 ligands across MSM-derived binding hotspots. Mean Gnina CNN affinity score (± SD across the 40 MSM-derived hotspot pockets) for the top 38 ranked compounds from a structure-based screen against Aβ42. Compounds (L1-L38) are ordered by decreasing mean CNN affinity along the x-axis, with all compounds shown exhibiting mean predicted affinity greater than 7.0. Compounds were obtained by first scoring the TargetMol isomeric library (1.66 million compounds) against the full Aβ42 sequence using Ligand Transformer, retaining the top-scoring subset for structure-based evaluation. Each candidate was then independently docked against all 40 hotspot pockets, each pocket corresponding to a representative frame of a high-equilibrium-probability microstate from the Aβ42-G5 MSM. Compounds were ranked by their mean Gnina CNN affinity across all pockets. Application of Lipinski’s rule of five and predicted aqueous solubility thresholds reduced this set to nineteen compounds advanced for prospective experimental evaluation. The narrow spread of mean scores and small inter-pocket standard deviations indicate that the top-ranked candidates engage the MSM-derived hotspots consistently across the Aβ42 conformational ensemble.

Application of Lipinski’s rule of five and predicted aqueous solubility filters to the top 200 narrowed the set to 19 compounds, which were advanced to experimental validation. These 19 candidates collectively cover multiple chemotypes and are predicted to engage a distributed subset of the 40 hotspot conformations rather than clustering on a single dominant site, in line with the protein-defined, multi-hotspot picture developed in the preceding sections. Experimental affinity measurements on these compounds will provide a prospective test of whether the binding-path framework can identify Aβ42 ligands improving on previously reported binders such as G5.

## Discussion

The central question motivating this work was whether binding pathways on disordered proteins are primarily defined by the ligand or by the protein. The data presented here support the second alternative. Across two disease-relevant disordered systems (Aβ42 and the C-terminal region of α-synuclein), and a chemically diverse panel of small molecules, the dominant residue-contact features identified by Markov state modelling are largely conserved. Pairwise enrichment factors of 15-45 for the top-ranked contact triplets, AUC values typically above 0.75, and recovery of many dominant features within the first 20-30% of the ranked list together indicate that ligands of different chemistry sample a common interaction network. This conservation extends beyond ligand identity to environmental perturbation: for Aβ42-G5, the ranked features are preserved across changes in pH and temperature, suggesting that the underlying transition network is a robust property of the peptide conformational ensemble rather than an artefact of a particular simulation condition.

The urea control is relevant to the interpretation of these results. Without it, the high enrichment seen across specific binders could in principle reflect a floor effect of the analysis itself, as any molecule diffusing on a peptide surface might produce apparently conserved contact features, in which case the conservation would be an artefact of the method rather than a property of the protein. The analysis can therefore distinguish a non-specific binder from specific ones, which means the strong overlap observed among the designed ligands is a genuine signal rather than a methodological baseline. The conserved network reflects protein-defined recognition features, and ligands modulate the weights and residence times along it without substantially redrawing its topology.

The ensemble docking analysis provides an orthogonal computational test of the same idea. When candidate binding regions identified from the holo ensembles are scored against a large, chemically diverse library, the same regions tend to recur as favourable docking sites, whereas low-scoring regions remain comparatively unfavourable. The signal therefore does not depend only on the chemistry of a single ligand or on the optimisation of a single ligand-pose pair. Rather, the docking analysis is consistent with the presence of recurrent ligand-accessible regions within the peptide conformational ensemble. Together with the MSM analysis, this supports a picture in which binding to these disordered systems is governed by persistent interaction motifs rather than by ligand-induced formation of entirely new pockets.

These observations have a direct translational consequence, which we explored prospectively for Aβ42. If binding hotspots are substantially protein-defined, they can be mapped from the conformational ensemble of the target and then used to prioritise chemistry, rather than re-derived independently for every chemical series. The pipeline implemented here combines MSM-derived hotspots with sequence-based Ligand-Transformer scoring of 1.66 million TargetMol compounds and structure-based Gnina docking against the hotspot ensemble, followed by drug-likeness and solubility filters. This pipeline follows directly from our model, as Ligand-Transformer ranks compounds against the full Aβ42 sequence, and docking tests them against the recurrent surface features identified by the MSM. The 19 compounds nominated for experimental evaluation will provide a prospective test of whether the framework can identify new Aβ42 ligands, and whether any improve on previously reported binders such as G5.

More broadly, this work suggests a reframing of the design problem for disordered targets. The conventional pocket-fitting paradigm asks how a ligand can be made complementary to a defined cavity. For disordered proteins, where no such cavity persists, the analogous question is how a ligand can be made compatible with the dynamic pathway network generated by the conformational ensemble. Transition matrices and MSMs provide a practical way to map this network, and the conservation observed here suggests that such maps can be reused across related ligands.

Several scope and methodological limitations should be acknowledged. The most important concerns what is actually being compared: the enrichment analysis ranks residue-contact features rather than complete time-ordered trajectories or transition-path fluxes. The strongest conclusion supported by the data is therefore conservation of dominant contact motifs and local pathway features. Whether full pathway fluxes are conserved to the same extent remains to be tested. Beyond this, the systems studied here are short peptides or a peptide-length segment of α-synuclein, and whether the same degree of conservation holds for longer disordered proteins, or for disordered regions embedded in folded domains, is an open question. The MSMs are built from enhanced-sampling simulations whose accuracy depends on the underlying force field, the chosen collective variables, the contact definition, and the degree of sampling. The ensemble docking step inherits the limitations of pocket detection and scoring functions, and because the receptor conformations are drawn from holo ensembles, it should be viewed as computational support for the MSM analysis rather than as independent validation. Experimental testing of the prospective Aβ42 candidates, and extension of the analysis to additional disordered targets and larger ligand panels, will therefore be important to assess the generality of the conclusions.

These results indicate that disordered protein sequences shape preferred contact regions within their conformational ensembles, and that chemically diverse ligands sample these regions to different extents rather than defining their own. For disordered targets, druggability may therefore reside less in a persistent pocket than in a recurrent network of contact regions encoded by the sequence.

## Materials and Methods

### Disordered protein systems

Two disordered systems were analysed in this work. Aβ42 (42 residues; UniProt P05067, residues 672-713 of amyloid precursor protein) was simulated in complex with three ligands (G5, D4, D8) and under three environmental conditions (pH 5 at 278 K, pH 5 at 298 K, pH 7 at 278 K) with G5. The C-terminal region of α-synuclein (residues 121-140 of UniProt P37840) was analysed in complex with seven ligands (Lig_4, Lig_12, Lig_20, Lig_26, Lig_30, Lig_40, Lig_50) using previously published all-atom molecular dynamics trajectories.

### Aβ42 simulations

All-atom metainference-metadynamics simulations^27,28^ of Aβ42 in complex with their respective ligands were performed using GROMACS 2018.3^29^ in combination with PLUMED 2.6.0-dev^30^, with the a99SB-disp force field^31^ and the TIP4P-D water model^32^ for the protein and water, and SAGE (openff)^33^ for the small molecule ligands. pH-dependent protonation states of titratable residues in Aβ42 and G5 were assigned using PROPKA 3.4^34^ for each of the three Aβ42-G5 conditions (pH 5 / 278 K, pH 5 / 298 K, pH 7 / 278 K). Each system was energy-minimised using steepest descent until the maximum force fell below 1000 kJ mol^-1^ nm^-1^, then equilibrated in two stages: 500 ps in the NVT ensemble using the Bussi-Donadio-Parrinello (v-rescale) thermostat^35^ with a coupling time of 0.2 ps, and 500 ps in the NPT ensemble with the Berendsen barostat^36^ at 1 bar with a coupling time of 1.0 ps. Production simulations were carried out in the NPT ensemble using the Parrinello-Rahman barostat^37^ at 1 bar, at the system-specific target temperature (278 K for Aβ42-G5 at both pH 5 / 278 K and pH 7 / 278 K, 298 K for Aβ42-G5 at pH 5 / 298 K). Short-range electrostatic and van der Waals interactions used a Verlet cutoff scheme with a 1.2 nm neighbour-list cutoff.

### α-synuclein C-terminal trajectories

Trajectories of the C-terminal region of α-synuclein in complex with the seven small molecules studied here were taken from the previously published 60 µs unbiased atomistic molecular dynamics simulations^19^, performed on the Anton 2 supercomputer using the a99SB-disp force field^31^ and the TIP4P-D water model^32^. From the full-length α-synuclein trajectories, frames were extracted and the analysis was restricted to residues 121-140 of α-synuclein. No additional simulation was performed for these systems. Frame weights for these trajectories are uniform, since the underlying trajectories are unbiased.

### MSM construction

Markov state models of ligand binding were constructed independently for each protein-ligand-condition system, following a framework previously described^23^, at each frame of the trajectory, all residues within a heavy-atom contact distance of 3.0 Å of any ligand atom were identified. From these contacts, the most populated quintuplet of simultaneously contacting residues at each frame was selected as the microstate assignment for that frame. Quintuplets were chosen as the MSM state definition on the basis of a Chapman-Kolmogorov validation^23^, and re-verified for the new systems analysed here. Transitions between microstates were counted at a lag time of 10 ns. An explicit unbound state was introduced at the beginning and end of each trajectory to avoid the creation of artificially absorbing states. Duplicate quintuplet states were filtered, and the resulting transition count matrix was row-normalised to yield the transition probability matrix T.

The equilibrium distribution π was obtained by eigendecomposition of T, with π corresponding to the left eigenvector associated with the eigenvalue λ₀ = 1. State-specific dissociation constants were calculated as K_D,i_ = π_u_/π_i_, where π_u_ is the equilibrium population of the unbound state and π_i_ is the equilibrium population of microstate i. Global dissociation constants were computed as K_D_ = (π_u_ / Σ_i_ π_i_) · (1 / N_A_V), where V is the simulation box volume and N_A_ is Avogadro’s number. Microstates were aggregated into macrostates by Louvain community detection^38^ applied to the microstate transition graph^23^. Macrostate equilibrium populations were obtained by summing microstate populations within each community, and degree centrality was computed on the coarse-grained macrostate graph using NetworkX^39^.

### Cross-system pathway comparison (enrichment factor and AUC analyses)

To assess the conservation of dominant binding pathways across ligands and conditions, ranked pathway ensembles were compared pairwise using enrichment factor (EF) and area-under-the-curve (AUC) analyses on contact. For each system, contact triplets were ranked by their equilibrium probability, defined as the marginal probability that a given residue triplet appears in any microstate of the equilibrium distribution. The total number of distinct triplets N was system-specific. The enrichment factor at cutoff k was calculated as

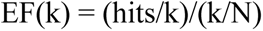

where hits is the number of triplets that appear in the top k of both ranked lists in a pairwise comparison. EF values were computed at top-k cutoffs of 20, 50, 100, 200, and 400. The expected value of EF under random ranking is 1. Cumulative recovery curves were constructed by ranking the triplets of system A from highest to lowest equilibrium probability and recording the fraction of the top 30 triplets of system B recovered as the ranked list of A was scanned. AUC values were computed by trapezoidal integration, with AUC = 0.5 corresponding to random expectation and AUC = 1.0 to perfect ranking agreement. To estimate statistical confidence, 95% confidence intervals on EF and AUC values were obtained by bootstrapping over 1000 resamples of the trajectory frames, with each resample independently used to construct a transition matrix and rank triplets.

### Ensemble docking

For each holo trajectory, 100 frames were selected as receptor templates. For each frame, the bound ligand was removed and candidate binding pockets were identified using Fpocket 4.0^40^; the top 5 pockets per frame, ranked by Fpocket druggability score, were retained. A chemically diverse small-molecule library was constructed from approximately 7 million compounds in the ZINC15 database^41^. Compounds were filtered for compliance with Lipinski’s rule of five^42^ and clustered with the Butina algorithm^43^ using Morgan fingerprints (radius 2, 2048 bits) generated with RDKit^44^ and a Tanimoto similarity cutoff of 0.4]. Ligands were prepared using RDKit and Meeko^45^, including protonation at pH 7.4, hydrogen addition, 3D coordinate generation, and conversion to PDBQT format. Receptor PDBQT files were prepared using AutoDockTools^46^. Docking was performed using AutoDock Vina 1.2^47^ centred on the pocket centroid as identified by Fpocket. To evaluate pocket-level binding propensity, the distribution of Vina docking scores across the chemical library was computed for each pocket.

### Recovery of binding hotspots

To compare the ability of ensemble docking and Fpocket alone to identify productive binding regions, the true binding pocket was defined as the set of residues in contact with the bound ligand in the corresponding holo frame, using the 3 Å heavy-atom cutoff used elsewhere in this work. Ensemble docking predictions were defined as the top pocket by mean Vina score across the chemical library; Fpocket predictions were defined as the top pocket by Fpocket druggability score on the same frame.

### Prospective ligand screening for Aβ42

For the prospective Aβ42 screen, the TargetMol isomeric compound library (1.66 million compounds) was scored against the wild-type Aβ42 amino acid sequence using Ligand-Transformer^24^, a sequence-based transformer model that predicts protein-ligand binding affinity from protein sequence and ligand SMILES without requiring a three-dimensional receptor structure. The top 20,000 compounds ranked by predicted pK_D_ were retained for structure-based docking. Receptor structures were extracted from the Aβ42-G5 MSM (pH 7 / 278 K condition). The top 40 microstates by equilibrium probability were identified; for each microstate, the representative frame was taken as the midpoint of the longest continuous block of trajectory frames assigned to that state. The G5 ligand was removed from each representative frame to generate 40 apo receptor structures. The 20,000 retained candidate ligands were converted from SMILES to PDBQT format using Open Babel 3.1.1^48^ with the ‘--gen3d’ flag and Gasteiger partial charges, and stereochemistry was preserved during 3D coordinate generation. Each compound was docked independently into each of the 40 hotspot pockets using Gnina 1.0^25^ with a 20 × 20 × 20 Å. Compounds were ranked by their mean Gnina CNN affinity score across all 40 pockets.

## Supplementary Information

**Figure S1.**
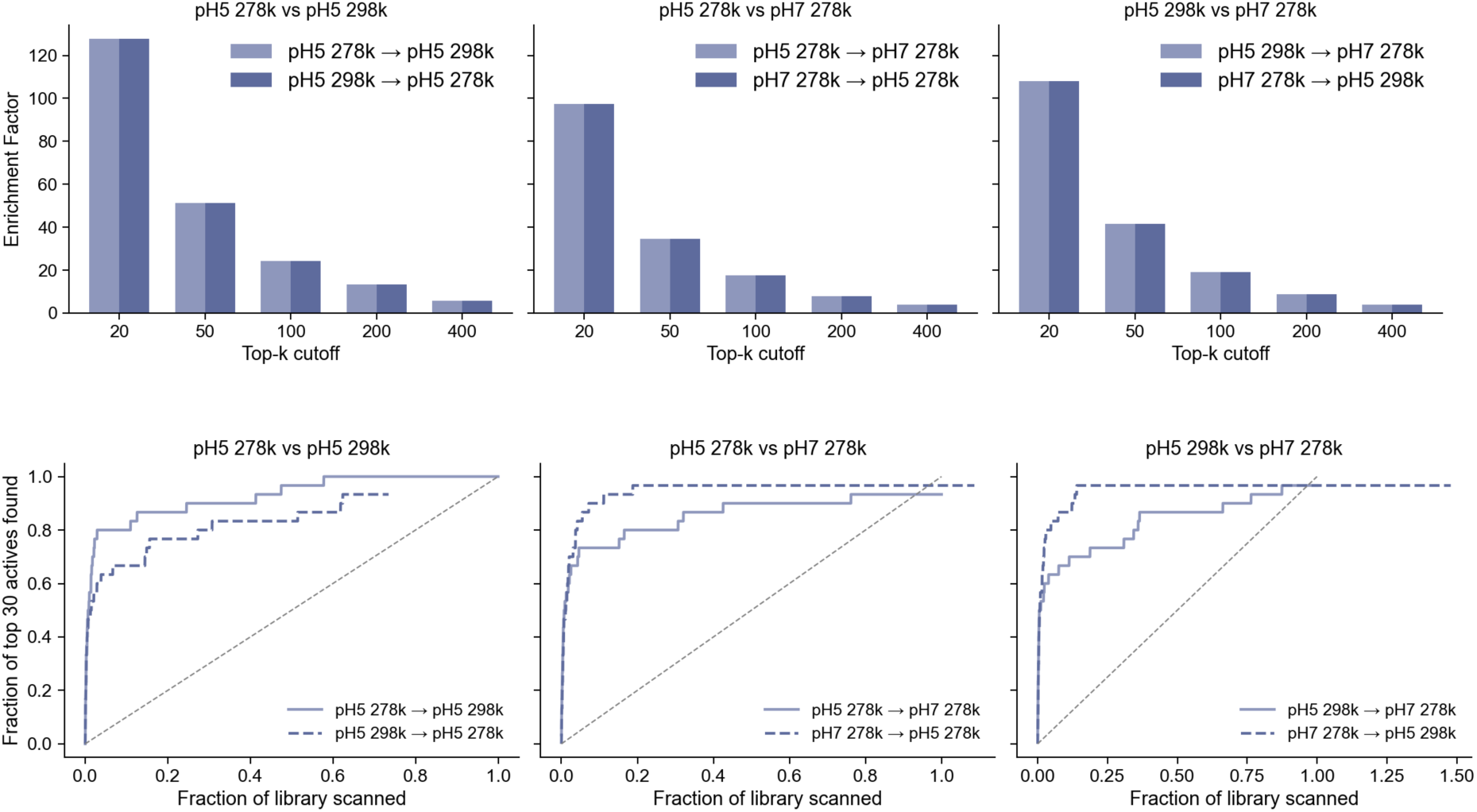
Conservation of binding pathways across pH and temperature for the Aβ42–G5 system. Cross-condition comparison of ranked binding pathways for Aβ42 in complex with G5 simulated under three physiological conditions: pH 5 at 278 K, pH 5 at 298 K, and pH 7 at 278 K. (Top row) Pairwise enrichment factors for overlap of top-ranked contact triplets across condition pairs (top-k cutoffs of 20, 50, 100, 200, and 400), with directional comparisons shown in light and dark blue. Enrichment factors exceed 100 at the smallest cutoffs and decay smoothly with increasing k, indicating strong recovery of the highest-ranked pathways across all condition pairs. (Bottom row) Cumulative recovery curves showing the fraction of top 30 reference states recovered as the ranked list of the comparison condition is scanned from highest to lowest probability. The diagonal grey line indicates random expectation. Across all three pairwise comparisons (pH 5 / 278 K vs. pH 5 / 298 K; pH 5 / 278 K vs. pH 7 / 278 K; pH 5 / 298 K vs. pH 7 / 278 K), recovery rises rapidly above the random baseline, with the majority of reference states recovered within the first 20-30% of the ranked list. Together, these results indicate that the dominant Aβ42-G5 binding pathways are conserved with respect to both pH and temperature within the physiologically relevant range examined.

**Figure S2.**
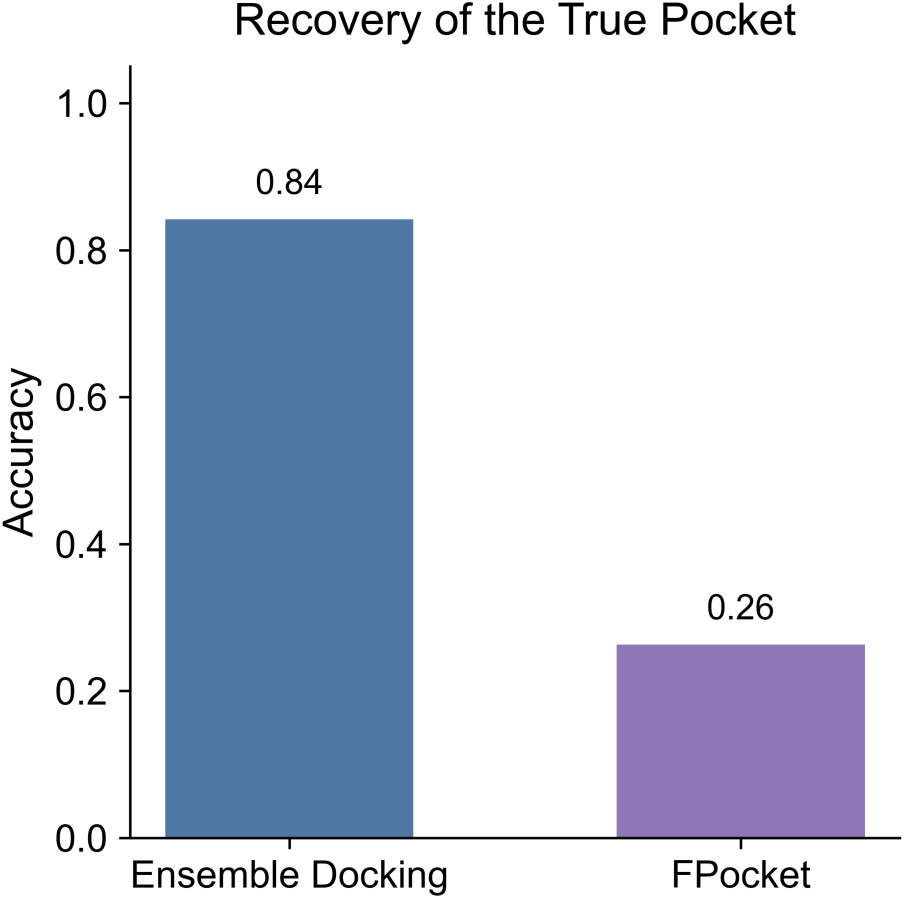
Recovery of the true binding pocket by ensemble docking and by Fpocket. Accuracy of binding pocket identification, defined as the fraction of holo trajectory frames in which the predicted top-ranked pocket overlaps with the experimentally observed ligand-binding region. Ensemble docking (blue), in which Fpocket-identified candidate pockets are evaluated by AutoDock Vina across multiple representative frames and ranked by docking score, recovers the true pocket in 89% of cases. Fpocket alone (purple), in which pockets are ranked by their intrinsic geometric and physicochemical scores without docking, recovers the true pocket in 26% of cases. The substantial improvement obtained by ensemble docking demonstrates that geometric pocket detection on a single frame is insufficient for disordered systems, and that combining ensemble sampling with ligand-based scoring is required to robustly identify productive binding regions on the conformational ensemble of an disordered protein.

**Figure S3.**
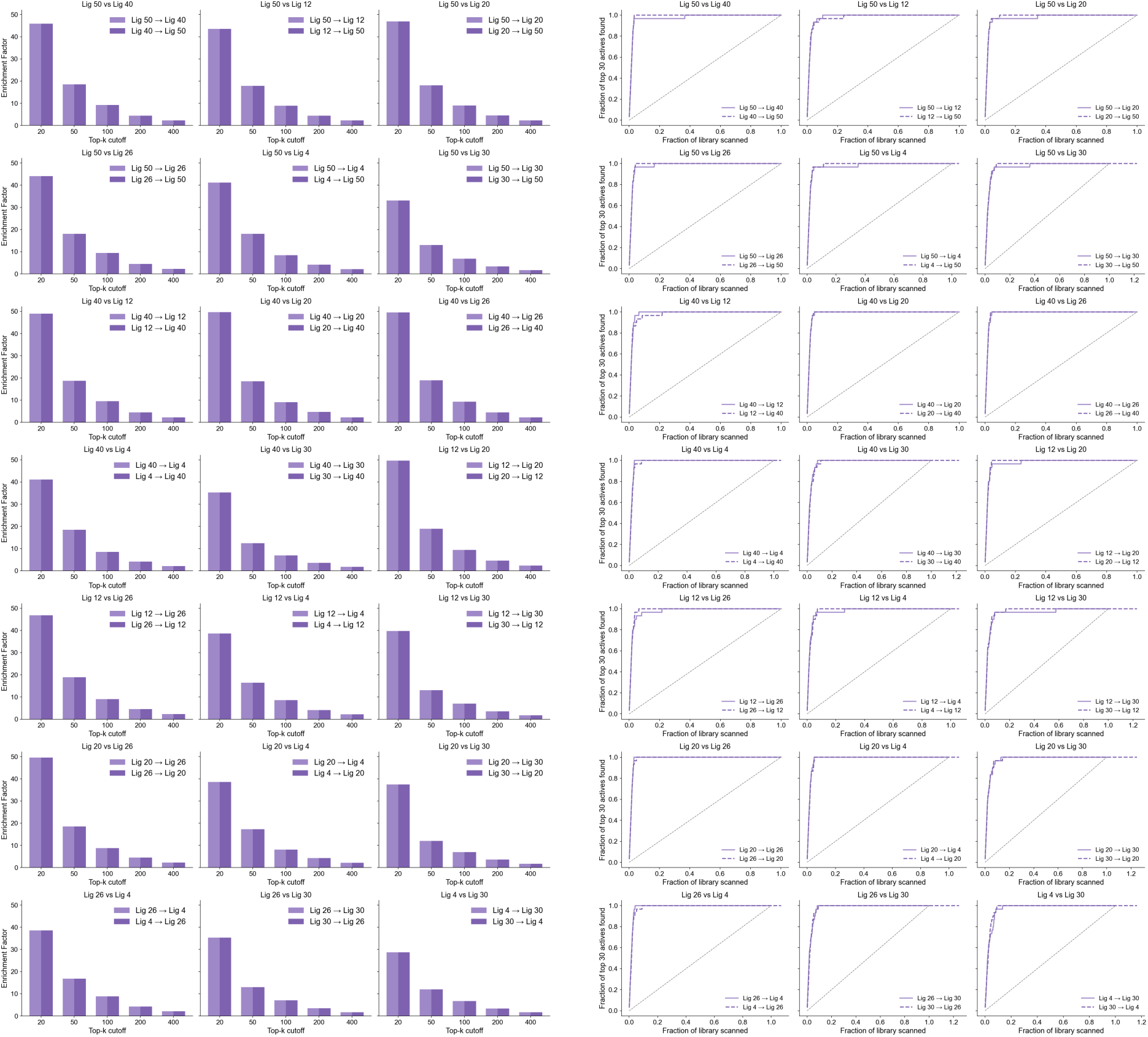
Pairwise consistency of ranked binding pathways across the seven α-synuclein C-terminal ligands. Cross-ligand comparison of binding-path ensembles for the seven small molecules (Lig 4, Lig 12, Lig 20, Lig 26, Lig 30, Lig 40, and Lig 50) studied in complex with the C-terminal region of α-synuclein. (Left block) Pairwise enrichment factors for overlap of top-ranked contact triplets, computed for each ordered ligand pair at top-k cutoffs of 20, 50, 100, 200, and 400. Each subpanel reports the two directional comparisons (e.g. Lig 50 → Lig 40 and Lig 40 → to Lig 50) in light and dark purple, and enrichment factors are well above unity at small cutoffs across nearly all ligand pairs, decaying smoothly with increasing k. (Right block) Corresponding cumulative recovery curves showing the fraction of top 30 reference states recovered as the ranked list of the comparison ligand is scanned. The diagonal grey line indicates random expectation. Across the full set of pairwise comparisons, recovery curves rise rapidly above the random baseline, with reference states typically recovered within the first 20-30% of the ranked list. Together, these analyses demonstrate that the conservation of dominant binding pathways observed for Aβ42 extends to the α-synuclein C-terminal system, with chemically diverse ligands consistently sampling the same high-probability residue-contact pathways.

