## Supplemental Information for "Does the sequence of a disordered protein encode small molecule binding paths?"

### Supplementary Information

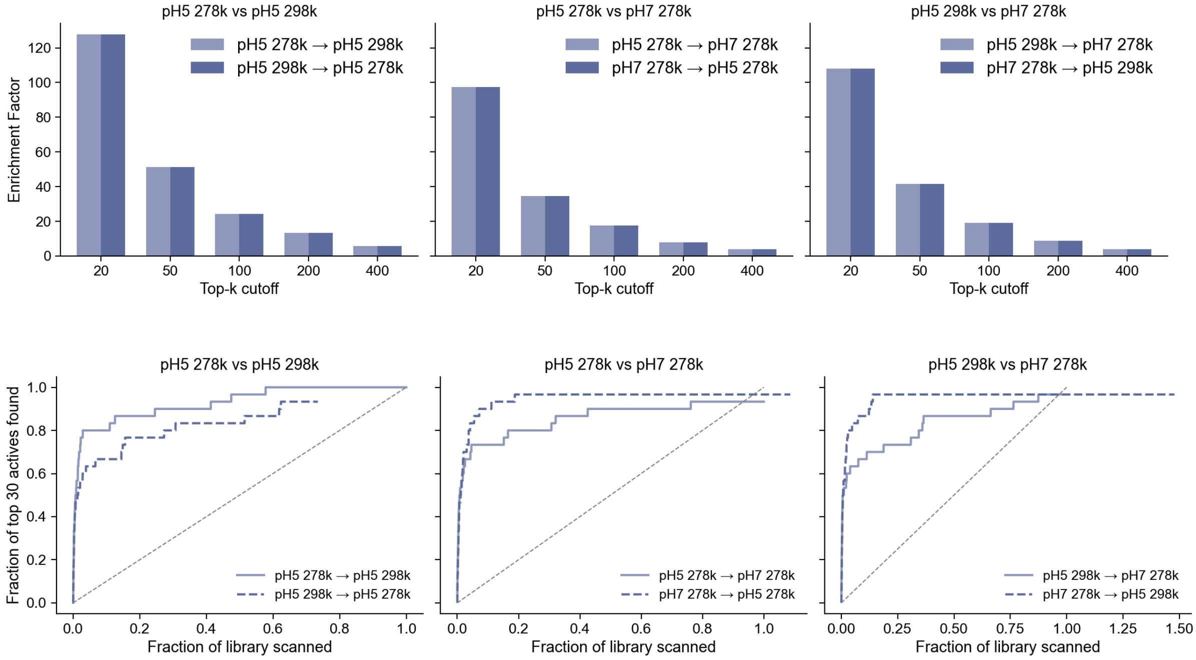


**Figure S1. Conservation of binding pathways across pH and temperature for the Aβ42–G5 system.** Cross-condition comparison of ranked binding pathways for Aβ42 in complex with G5 simulated under three physiological conditions: pH 5 at 278 K, pH 5 at 298 K, and pH 7 at 278 K. (Top row) Pairwise enrichment factors for overlap of top-ranked contact triplets across condition pairs (top-k cutoffs of 20, 50, 100, 200, and 400), with directional comparisons shown in light and dark blue. Enrichment factors exceed 100 at the smallest cutoffs and decay smoothly with increasing k, indicating strong recovery of the highest-ranked pathways across all condition pairs. (Bottom row) Cumulative recovery curves showing the fraction of top 30 reference states recovered as the ranked list of the comparison condition is scanned from highest to lowest probability. The diagonal grey line indicates random expectation. Across all three pairwise comparisons (pH 5 / 278 K vs. pH 5 / 298 K; pH 5 / 278 K vs. pH 7 / 278 K; pH 5 / 298 K vs. pH 7 / 278 K), recovery rises rapidly above the random baseline, with the majority of reference states recovered within the first 20-30% of the ranked list. Together, these results indicate that the dominant Aβ42-G5 binding pathways are conserved with respect to both pH and temperature within the physiologically relevant range examined.

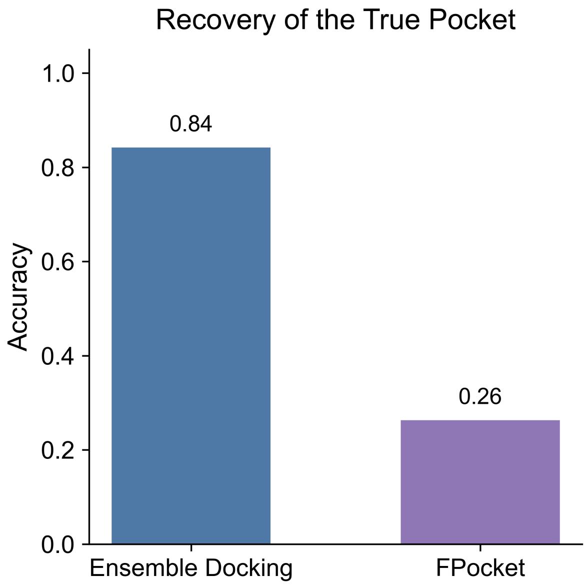


**Figure S2. Recovery of the true binding pocket by ensemble docking and by Fpocket.** Accuracy of binding pocket identification, defined as the fraction of holo trajectory frames in which the predicted top-ranked pocket overlaps with the experimentally observed ligand-binding region. Ensemble docking (blue), in which Fpocket-identified candidate pockets are evaluated by AutoDock Vina across multiple representative frames and ranked by docking score, recovers the true pocket in 89% of cases. Fpocket alone (purple), in which pockets are ranked by their intrinsic geometric and physicochemical scores without docking, recovers the true pocket in 26% of cases. The substantial improvement obtained by ensemble docking demonstrates that geometric pocket detection on a single frame is insufficient for disordered systems, and that combining ensemble sampling with ligand-based scoring is required to robustly identify productive binding regions on the conformational ensemble of an disordered protein.

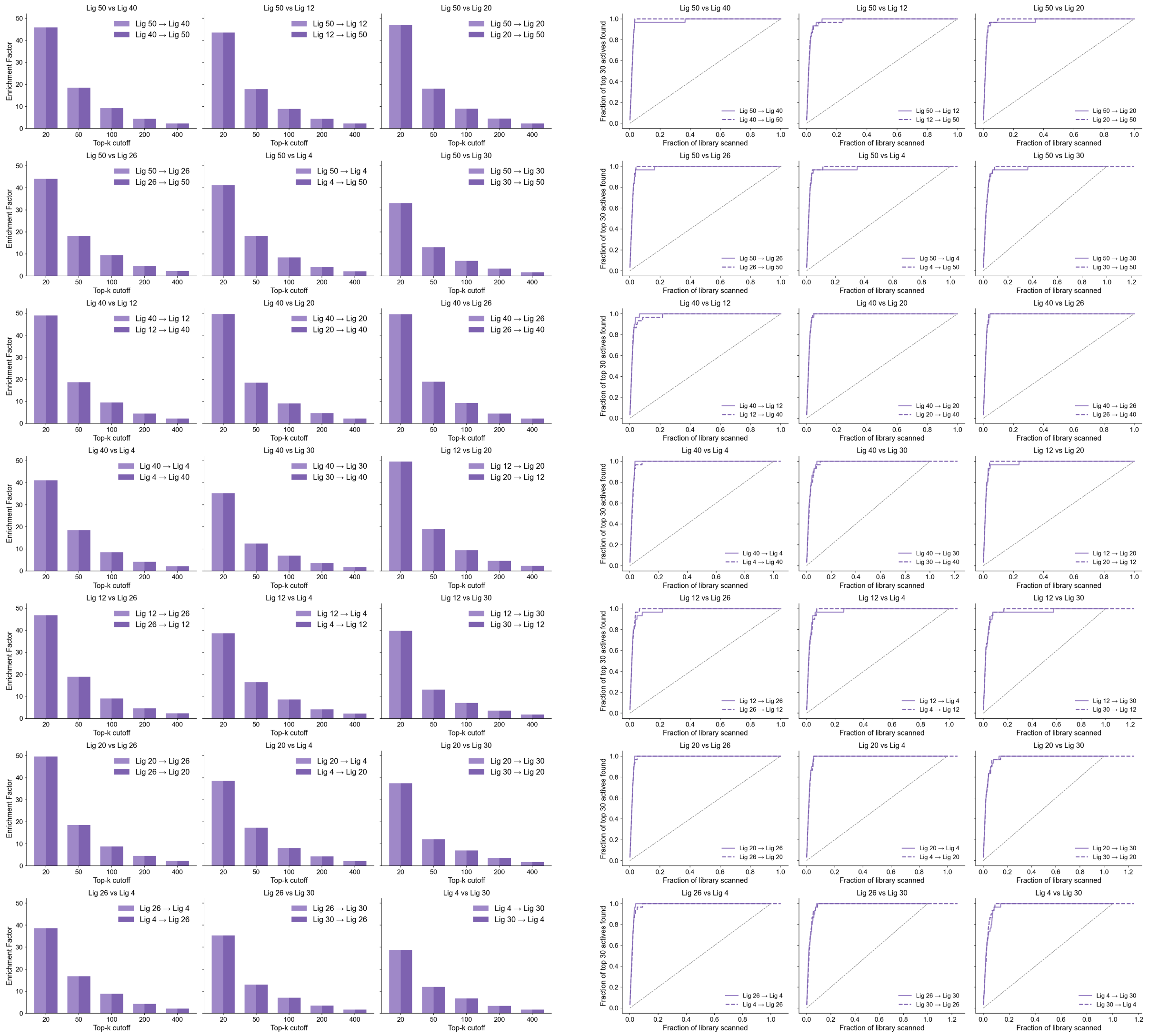


**Figure S3. Pairwise consistency of ranked binding pathways across the seven α-synuclein C-terminal ligands.** Cross-ligand comparison of binding-path ensembles for the seven small molecules (Lig 4, Lig 12, Lig 20, Lig 26, Lig 30, Lig 40, and Lig 50) studied in complex with the C-terminal region of α-synuclein. (Left block) Pairwise enrichment factors for overlap of top-ranked contact triplets, computed for each ordered ligand pair at top-k cutoffs of 20, 50, 100, 200, and 400. Each subpanel reports the two directional comparisons (e.g. Lig 50 → Lig 40 and Lig 40 → to Lig 50) in light and dark purple, and enrichment factors are well above unity at small cutoffs across nearly all ligand pairs, decaying smoothly with increasing k. (Right block) Corresponding cumulative recovery curves showing the fraction of top 30 reference states recovered as the ranked list of the comparison ligand is scanned. The diagonal grey line indicates random expectation. Across the full set of pairwise comparisons, recovery curves rise rapidly above the random baseline, with reference states typically recovered within the first 20-30% of the ranked list. Together, these analyses demonstrate that the conservation of dominant binding pathways observed for Aβ42 extends to the α-synuclein C-terminal system, with chemically diverse ligands consistently sampling the same high-probability residue-contact pathways.
